# Role of the imprinted allele of the p57^Kip2^ gene in mouse neocortical development

**DOI:** 10.1101/737692

**Authors:** Yui Imaizumi, Shohei Furutachi, Tomoyuki Watanabe, Hiroaki Miya, Daichi Kawaguchi, Yukiko Gotoh

**Affiliations:** Graduate School of Pharmaceutical Sciences, The University of Tokyo, Tokyo 113-0033, Japan; Sainsbury Wellcome Centre for Neural Circuits and Behaviour, University College London, London W1T 4JG, UK

**Author notes:** Correspondence and requests for materials should be addressed to Y.G.

## Abstract

Imprinted genes are expressed from only one allele in a parent of origin–specific manner. The cyclin-dependent kinase inhibitor p57^Kip2^ (p57) is encoded by an imprinted gene, with the paternal allele being silenced. The possible expression and function of the paternal allele of p57 have remained little studied, however. We now show that the paternal allele of the p57 gene is expressed at a low level in the developing mouse neocortex. Surprisingly, the central nervous system-specific conditional deletion of the paternal allele (*pat* cKO) at the p57 locus resulted in a marked reduction in brain size. Furthermore, *pat* cKO gradually reduced the number of neural stem-progenitor cells (NPCs) during neocortical development, and thus reduced the number of upper-layer neurons, which were derived from late-stage NPCs. Our results thus show that the paternal allele of the p57 locus plays a key role in maintenance of NPCs during neocortical development.

## Introduction

Genomic imprinting refers to an epigenetic process that results in the inactivation of one of the two alleles of a gene in a parent of origin–dependent manner^1-3^. It is achieved mainly by allele-specific DNA methylation at a subset of CpG islands, known as imprinting control regions (ICRs), during early developmental stages^4^. Imprinted genes play essential roles in development, homeostasis, and behavior in mammals ^5,6^. Changes at imprinted gene loci in humans are associated with diseases such as Beckwith-Wiedemann syndrome, Prader-Willi syndrome, and Angelman syndrome^7,8^, many of which are characterized by altered growth and mental disorders^6^. Although canonical genomic imprinting has been thought to result in the complete silencing of one allele of a gene, recent studies have shown that silencing of some imprinted genes appears to be incomplete or reversed to various extents in the brain^9^. For instance, derepressed expression of the imprinted alleles of *Igf2* and *Dlk1* contributes to the regulation of adult neural stem cells in mice^10,11^. However, the functions of the imprinted alleles of other genes remain mostly obscure.

The p57^Kip2^ (p57) gene is imprinted, with the maternal allele being expressed, and is located in the distal region of mouse chromosome 7 and human chromosome 11p15^12,13^. In mice, DNA methylation of two ICRs, KvDMR^14-16^ and ICG5^17^, has been suggested to suppress expression of the paternal allele of the p57 gene (*Cdkn1c*). The p57 protein is a cyclin-dependent kinase inhibitor (CKI)^18,19^ that is highly expressed in neural and skeletomuscular tissues^20,21^ during embryonic development. In humans, changes at the p57 gene locus are associated with Beckwith-Wiedemann syndrome, features of which include excessive growth and an increased risk of childhood cancer^22,23^. Gene knockout (KO) studies have implicated p57 in regulation of fetal growth and placental development^20,24,25^. In the central nervous system (CNS), p57 plays a key role in regulation of the proliferation and differentiation of embryonic neural stem-progenitor cells (NPCs) and adult neural stem cells^26-33^. CNS-specific conditional KO of the maternal *p57* allele resulted in the induction of cell death in a manner dependent on the transcriptional regulators E2F1 and p53, thinning of the neocortex, and pronounced hydrocephalus in mice, with the latter effects possibly reflecting a function of p57 in the subcommissural organ (SCO)^30^.

The paternal allele of the p57 gene has been thought to be completely silenced, given that conventional KO of the maternal allele appeared to result in elimination of p57 mRNA and protein^20^ and that a reporter for the paternal allele did not show any expression during normal development unless challenged by stress^34^. Indeed, the phenotype of conventional maternal p57 KO mice appears essentially identical to that of conventional null p57 KO mice^20,24,32^, and paternal p57 KO mice have been found to manifest no distinct phenotype^24^. These observations have thus suggested that only the maternal allele of the p57 gene locus is indispensable. However, it has remained possible that the paternal allele of this locus is transcribed at a low level, and the necessity of the paternal allele for development has remained poorly explored.

Here we examined the expression and functional importance of the paternal *p57* allele in the developing mouse brain. Reverse transcription (RT) and allele-specific quantitative polymerase chain reaction (qPCR) analysis revealed that the paternal allele is indeed expressed at a low level in embryonic NPCs. Unexpectedly, CNS-specific paternal p57 KO resulted in a substantial reduction in brain size, with the number of upper layer neurons in the neocortex showing a marked decrease. Consistent with this latter observation, the number of NPCs that give rise to upper layer neurons was also decreased in the mutant mice. Our findings thus uncover an essential role for the imprinted allele of the p57 gene in the developing nervous system.

## Results

### Expression of the paternal *p57* allele in neocortical NPCs and neurons during mouse embryonic development

We first examined whether expression of the paternal (imprinted) allele of the p57 gene could be detected in the developing brain with the use of chimeric mice derived from a cross between lines—C57BL/6J (BL6) and JF1/Ms (JF1)—that differ with regard to single nucleotide polymorphisms (SNPs) at this locus. We thus collected F_1_ hybrid embryos at embryonic day (E) 16 from a cross between BL6 male and JF1 female mice. Total RNA isolated from the embryonic neocortex was then subjected to RT and allele-specific qPCR analysis with primers designed to recognize individual SNPs (Fig. 1a). In contrast to previous results showing that the paternal allele of the p57 gene is completely silenced^20,34^, we detected expression of the paternal allele at a level corresponding to ∼1% to 2% of that of the maternal allele (Fig. 1b). We did not detect p57 mRNA in embryos from a cross between BL6 male and BL6 female mice with the use of JF1-specific primers or in those from a cross between JF1 male and JF1 female mice with the use of BL6-specific primers, confirming the specificity of the line-specific primers. We next deleted the paternal allele of the p57 gene in the CNS by crossing JF1 female mice heterozygous for a Cre recombinase transgene under the control of the *Nestin* enhancer (*Nestin-Cre*) with BL6 male mice harboring floxed alleles of the p57 gene (*p57*^fl/fl^) (Fig. 1c). Expression of the paternal *p57* allele in the neocortex was substantially reduced in the resulting conditional KO (cKO) embryos at E16 compared with control littermates lacking the *Nestin-Cre* transgene (Fig. 1d). Given that the expression pattern of *Nestin-Cre*^35^ indicates that the floxed *p57* allele would have been deleted in NPCs of the CNS only after ∼E11, this result suggests that the paternal *p57* allele indeed contributed to the *p57* transcripts detected with BL6-specific primers in these embryos. We then examined expression of the paternal *p57* allele in NPCs and neurons isolated from the neocortex of E16 embryos derived from a cross between BL6 male and JF1 female mice. These cells were isolated as CD133^+^CD24^−^ and CD133^−^CD24^+^ fractions, respectively, by fluorescence-activated cell sorting (FACS), and both were found to express the paternal *p57* allele (Fig. 1b). Together, these results thus indicated that the paternal allele of the p57 gene is transcribed, albeit at a low level, in the developing neocortex.

**Figure 1.**
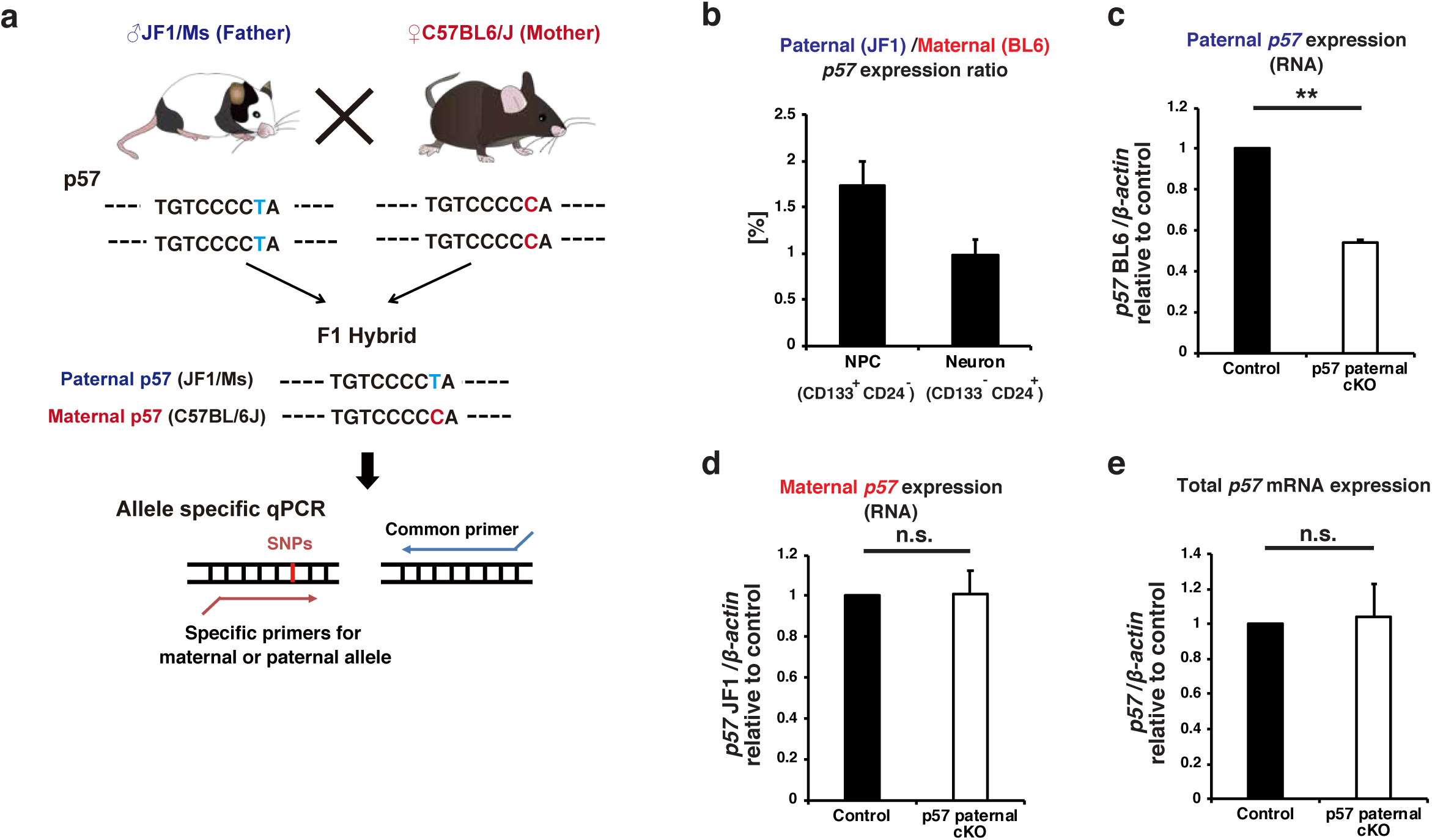
Detection of *p57* mRNA expression from the paternal allele in the embryonic neocortex. **(a)** Scheme of allele specific qPCR of p57. F1 hybrid embryos of C57BL/6J (BL6) female and JF1/MS (JF1) male parents have SNPs derived from their parents. qPCR was performed by the use of specific primers for each allele. **(b)** The ratio of paternal to maternal *p57* mRNA expression in NPCs and in neurons was determined by allele specific qPCR under the hybrid (JF1 and BL6) genetic background at E16. FACS was used to isolate NPCs (CD133^+^ CD24^−^ population) and neurons (CD133^−^ CD24^+^ population). **(c)(d)** Allele specific qPCR of *p57* mRNA in the neocortex isolated from control (*p57* pat flox (BL6)/+(JF1)) and hybrid paternal p57 cKO (Nestin-Cre; *p57* ^pat flox(BL6)/+(JF1)^) mice at E16. *p57* mRNA expression from each allele was normalized to β*-actin*. Data are expressed relative to the corresponding value for control mice. **(e)** qPCR of total p57 mRNA expression in the neocortex isolated from control and paternal p57 cKO mice in the BL/6J background at P0. *p57* mRNA expression was normalized to β*-actin*. Data are expressed relative to the corresponding value for control mice. Data are mean+s.e.m from three independent experiments. Paired two-tailed Student’s t-test; **\*\**P***<0.01. n.s., not significant.

We investigated whether deletion of the paternal *p57* allele might affect expression of the maternal allele. However, the amounts of maternal or total p57 mRNA in the neocortex did not differ significantly among E16 embryos derived from a cross between BL6 male *p57*^fl/fl^ and JF1 female *Nestin-Cre*^+/–^ mice (Fig. 1d and 1e). These results thus suggested that the paternal *p57* allele does not substantially affect the level of p57 expression from the maternal allele.

### CNS-specific paternal p57 KO reduces the brain size and the number of upper layer neurons at postnatal stage

We next investigated whether the paternal *p57* allele plays a role in mouse brain development. We deleted the paternal allele in a CNS-specific manner by crossing BL6 *p57*^fl/fl^ male with BL6 *Nestin-Cre*^+/–^ female mice and found that such deletion resulted in a substantial reduction in brain size apparent at postnatal day (P) 60 (Fig. 2a). This finding was unexpected given the low level of expression of the paternal allele compared with the maternal allele. Immunohistofluorescence analysis of coronal sections revealed that this size reduction was apparent throughout the entire forebrain—including the neocortex, basal ganglia, thalamus, and septal nuclei—of the paternal cKO mice at P24 (Fig. 2b). The total section size of the paternal p57 cKO brain at the level of 0.38 to 1.18 mm relative to the bregma was only 72.37 ± 6.07% (mean ± s.e.m., *n* = 3 mice) of that for the control brain at P24. With regard to the neocortex, the mediolateral surface length at this level relative to the bregma was reduced to 81.44 ± 1.53% (mean ± s.e.m., *n* = 3 mice) for the paternal p57 cKO brain compared with the control brain at P24. The thickness of the primary somatosensory area (S1) of the paternal p57 cKO brain was also reduced to 71.33 ± 10.00% (mean ± s.e.m., *n* = 3 mice) of that for the control brain at P24.

**Figure 2.**
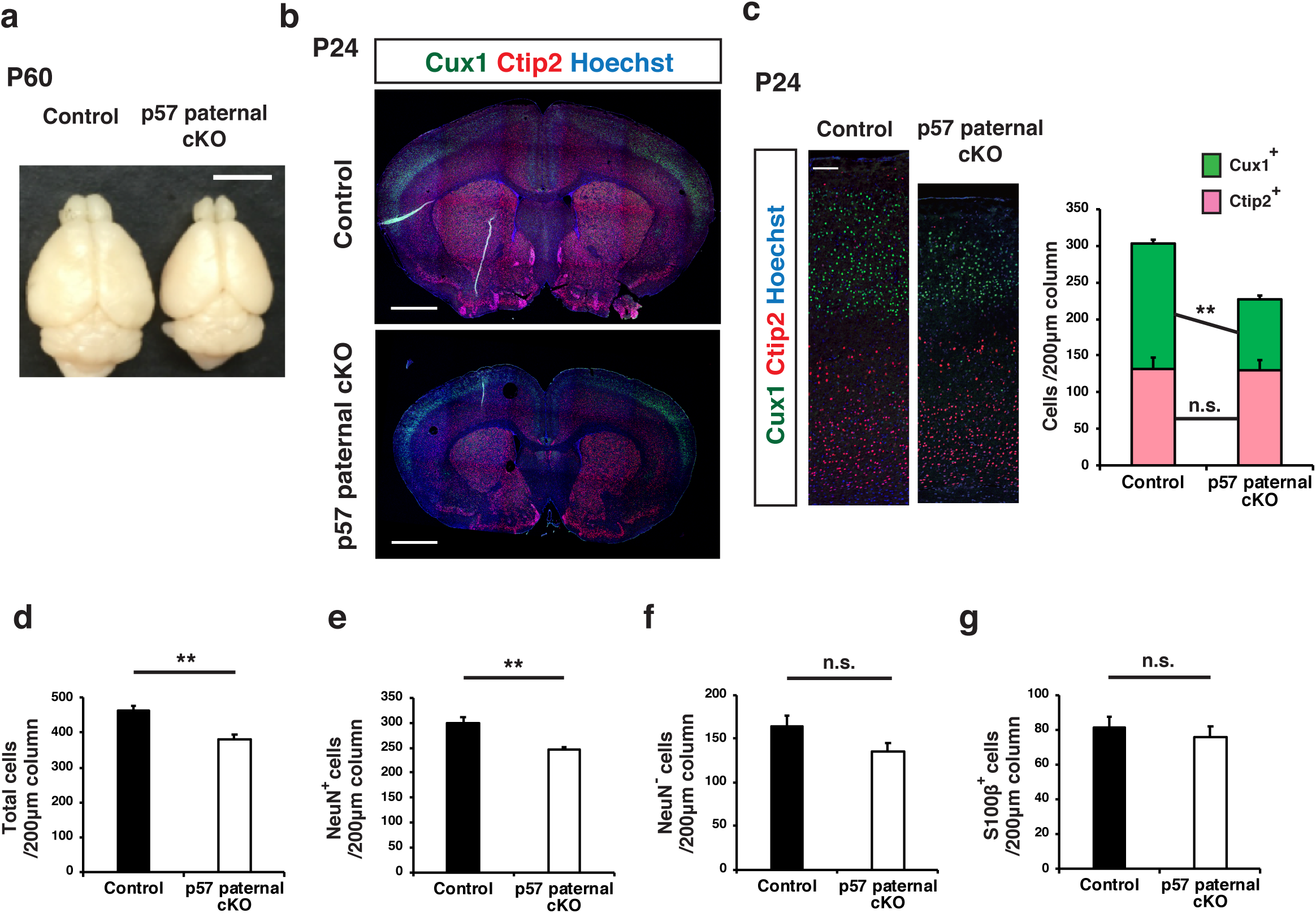
CNS-specific deletion of the p57 paternal allele reduced the brain size and the number of upper layer neurons at P24. **(a-g)** p57 paternal cKO (Nestin-Cre; *p57* ^pat flox/+^) and control (*p57* ^pat flox/+^) mice were sacrificed at P60 (**a**) and P24 (**b**-**g**). **(a)** Dorsal view of p57 paternal cKO and control brains. **(b)** Immunofluorescence staining of Cux1 (green) and Ctip2 (red) in coronal sections of p57 paternal cKO and control mice. Nuclei were stained with Hoechst (blue). **(c)** Higher magnifications of the neocortical somatosensory area in (**b**) (left panel). Quantitative analysis of cells positive for Cux1 and Ctip2 per area within 200 μm wide bins (right panel) (n = 3 mice for each genotype). **(d-g)** The number of total cells (Hoechst^+^) (**d**), neurons (NeuN^+^) (**e**), non-neuronal cells (NeuN^−^)(**f**) and glial cells (S100β^+^) (**g**) per 200 μm wide bins in the neocortical area were quantified (n = 4 mice for each genotype). Data are mean+s.e.m. Unpaired two-tailed Student’s t-test; \*\****P*** <0.01 n.s., not significant. Scale bars: 500 μm in (**a**); 250 μm in (**b**); 100 μm in (**c**).

The forebrain phenotype of the paternal p57 cKO mice appeared largely similar to that of maternal p57 cKO mice previously generated with the same genetic tools^30^, although the former was generally less pronounced and showed some distinct features. One such feature of the paternal cKO brain was the apparent absence of hydrocephalus, one of the most remarkable characteristics of the maternal p57 cKO brain^30^. The ventricles of the paternal p57 cKO brain were not as expanded as those of the control brain at P0 or P24 (Fig. 2b and 3a). The previous study of maternal p57 cKO mice concluded that the thinning of the neocortex also apparent in these mice was a result of the hydrocephalus caused by impaired development of the SCO^30^. Our results revealing a thin neocortex without apparent hydrocephalus in the paternal p57 cKO brain thus indicates that this neocortical phenotype is not due to hydrocephalus, at least not in these animals.

**Figure 3.**
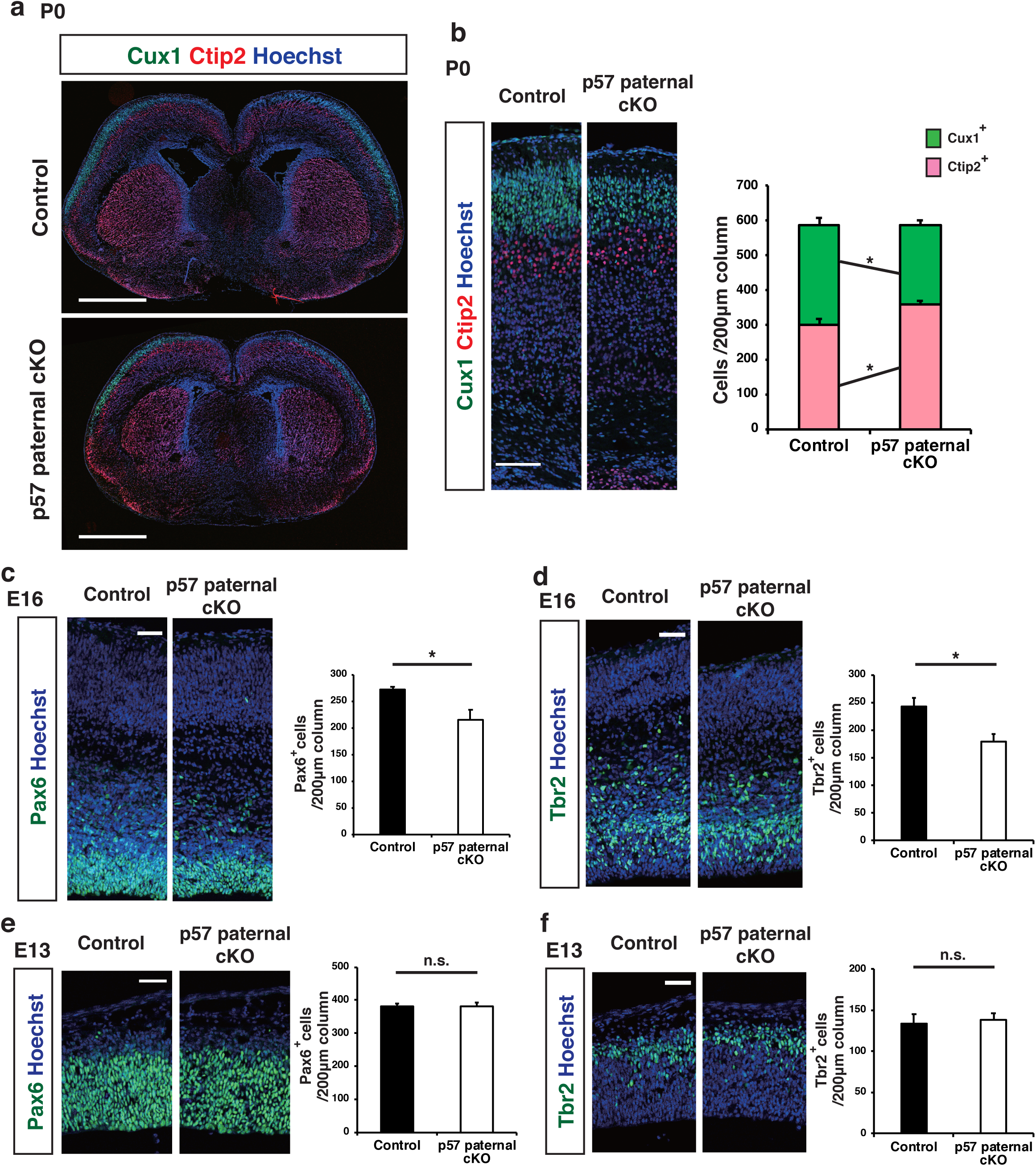
CNS-specific deletion of the p57 paternal allele reduced the number of upper layer neurons at P0 and progenitors at E16. **(a)** Immunofluorescence staining of Cux1 (green) and Ctip2 (red) in coronal sections of p57 paternal cKO and control mice at P0. Nuclei were stained with Hoechst (blue). **(b)** Higher magnifications of neocortical somatosensory area in **(a**) (left panel). Quantitative analysis of cells positive for Cux1 and Ctip2 per area within 200 μm wide bins (right panel) (n = 4 pups for each genotype). **(c-f)** Immunofluorescence staining of Pax6 (**c**,**e**) and Tbr2 (**d**,**f**) at E16 (**c**,**d**) and E13 (**e**,**f**), respectively. The cells positive for Pax6 and Tbr2 per area within 200 μm wide bins in the neocortical region were quantified (n=4 embryos for each genotype in **c**,**d** and n=3 embryos for each genotype in **e**,**f**). Data are mean+s.e.m. Unpaired two-tailed Student’s t-test; \****P*** <0.05. n.s., not significant. Scale bars: 500 μm in (**a**); 100 μm in (**b**); 50 μm in (**c**-**f**)

We next examined the neocortex of paternal p57 cKO mice at P24 in more detail. Consistent with the reduced thickness of the neocortex, the cell number in S1 was reduced in the paternal p57 cKO mice compared with control mice (Fig. 2d, Supplementary Fig. 1a). The number of NeuN^+^ neurons was reduced by paternal p57 cKO while the number of NeuN^−^ cells and S100β^+^ astrocytes was not (Fig. 2e, 2f and 2g). Among neurons, the number of Cux1^+^ upper layer neurons was significantly reduced whereas that of Ctip2^+^ deep layer neurons was not (Fig. 2c). This selective reduction in the number of upper layer neurons relative to deep layer neurons in the paternal p57 cKO neocortex at P24 was confirmed by the detection of a reduced thickness of the upper layers but not of the deep layers in S1 (Supplementary Fig. 1a). These findings contrast with those obtained by conventional KO of the maternal *p57* allele, which resulted in a selective increase in the number of deep layer neurons^32^. Our results thus suggested that the paternal *p57* allele might contribute to the genesis of specific neuronal subtypes.

### Paternal p57 cKO reduces the number of neocortical NPCs at late embryonic stages

We next examined when and how paternal p57 cKO results in a reduction in the number of upper layer neurons in the neocortex during development. We first investigated the neocortex of control and paternal p57 cKO mice at P0 and found that the reduction in the number of Cux1^+^ upper layer neurons was already apparent at this stage (Fig. 3a and 3b). Of note, the number of Ctip2^+^ deep layer neurons at this stage was actually increased by paternal p57 cKO (Fig. 3b). Consistent with these results, the thickness of the upper layers was reduced but that of the deep layers tended to increase in the paternal p57 cKO mice at P0 (Supplementary Fig. 1b).

To investigate the mechanism underlying the selective reduction in the number of upper layer neurons in the postnatal neocortex of paternal p57 cKO mice, we evaluated the production of deep layer and upper layer neurons at embryonic stages. We first examined the numbers of Pax6^+^ NPCs and Tbr2^+^ intermediate neuronal progenitors (INPs) in S1 at E13 and found no significant difference between control and paternal p57 cKO mice (Fig. 3e and 3f). However, paternal p57 cKO resulted in a significant reduction in the numbers of both these cell types at E16 (Fig. 3c and 3d). Consistent with these findings, the number of proliferating cells positive for Ki67 was also reduced in the neocortex of paternal p57 cKO mice at E16 but not at E13 (Fig. 4a and 4b). Given that most deep layer neurons and upper layer neurons in S1 are born before and after E13.5, respectively, the differences in the numbers of NPCs and INPs apparent at E16 but not at E13 may explain, at least in part, the selective reduction in the number of upper layer neurons induced by paternal p57 cKO. We also found that the number of apoptotic cells positive for the cleaved form of caspase-3 was increased among NPCs in the neocortex of paternal p57 cKO mice at both E13 and E16 (Fig. 4c and 4d). Related to apoptosis induction, we also detected increased expression of the gene for the CKI p21^Cip1^, a target of p53, in the neocortex of paternal p57 cKO mice at E16 (Fig. 4e). The increase in the extent of cell death might thus account for the reduction in the number of NPCs and subsequently that in INPs in the paternal p57 cKO neocortex. Together, these results indicate that the paternal *p57* allele promotes cell survival and plays an important role in the generation of appropriate numbers of NPCs, INPs, and cortical neurons during development.

**Figure 4.**
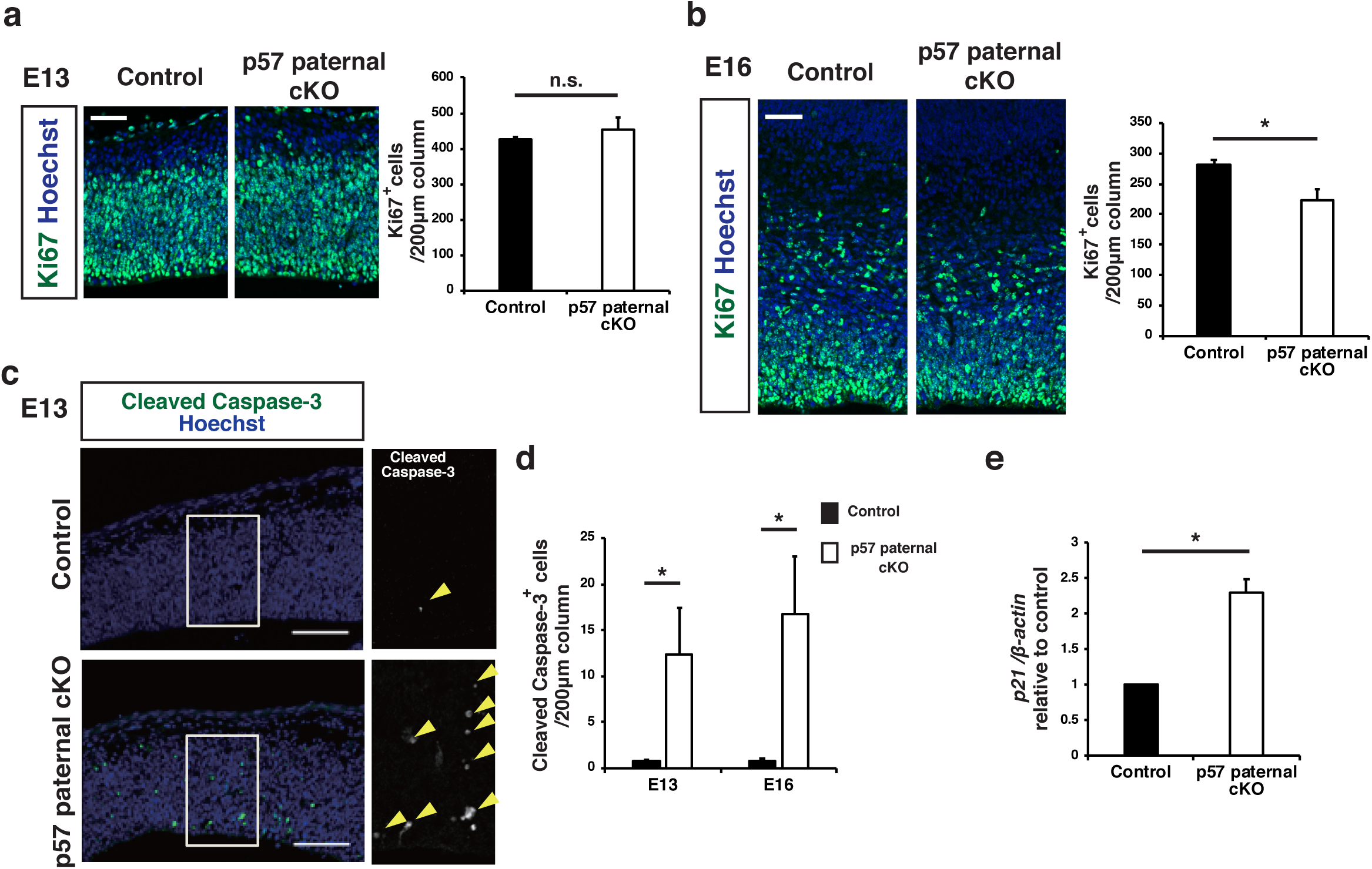
CNS-specific deletion of the p57 paternal allele reduced the number of proliferating progenitors at E16 and increased the number of apoptotic cells at E13 and E16. **(a-b)** Immunofluorescence staining of Ki67 at E13 (**a**) and E16 (**b**) (left panel). Quantitative analysis of cells positive for Ki67 per area within 200 μm wide bins (right panel) (n = 3-4 embryos for each genotype). Data are mean+s.e.m. Unpaired two-tailed Student’s t-test. **(c)** Immunostaining of cleaved caspase-3 at E13. The boxed regions in the left panels are shown at a higher magnification in the right panel. Yellow arrowheads indicate cleaved caspase-3 positive cells. **(d)** The number of cleaved caspase-3 positive cells per area within 500 μm wide bins were quantified at E13 and E16 (n=4-6 embryos for each genotype). Data are mean+s.e.m. Unpaired two-tailed Student’s t-test. **(e)** qPCR of *p21* mRNA expression in the neocortex isolated from control and paternal p57 cKO mice at P0. *p21* mRNA expression was normalized to β*-actin* (n=3 independent experiments). Data are mean+s.e.m.; expressed relative to the corresponding value for control mice. Paired two-tailed Student’s t-test. \****P***<0.05. n.s., not significant. Scale bars: 50 μm in (**a**) (**b**); 100 μm in (**c**)

## Discussion

We have here detected a low level of expression of the paternal *p57* allele in the developing mouse neocortex, in contrast to previous findings that the paternal allele is completely imprinted and silenced^20,34^. Immunohistochemical analysis thus previously showed that the amount of p57 protein was reduced to an undetectable level in the brain by deletion of the maternal *p57* allele^34^, and reporter mice that allow monitoring of the expression of the paternal allele on the basis of the activity of firefly luciferase encoded by a construct knocked-in at this locus did not show a luminescence signal under normal dietary conditions^34^. However, in contrast to the allele-specific qPCR analysis adopted in our study, expression of the paternal *p57* allele might have been too low compared with that of the maternal allele to have been detected by these previous methods.

Although expression of the paternal *p57* allele in the brain was <2% of that of the maternal allele at E16, we found that paternal p57 cKO resulted in prominent changes to the brain including thinning of the neocortex. Such thinning previously observed in maternal p57 cKO mice was suggested to be a result of defects in the SCO^30^. However, the neocortical thinning detected in our paternal p57 cKO mice appears to be independent of any effect on the SCO, given the absence of hydrocephalus. Instead, our results suggest that increased cell death and a consequent reduction in the number of neocortical NPCs at the late stage of development give rise to a reduced level of neurogenesis and thinning of the neocortex in paternal p57 cKO mice. Our results have thus revealed an essential role for the paternal *p57* allele in control of the survival of neocortical NPCs and the genesis of neurons—in particular, upper layer neurons produced at the late stage of development.

How the paternal *p57* allele fulfills this role despite its low expression level remains unknown. We measured expression of the paternal allele in populations of NPCs and neurons, so it is possible that the allele is expressed at similar or even higher levels relative to the maternal allele in a subpopulation of NPCs in which it plays a protective role. Alternatively, the paternal and maternal alleles might confer expression of different mRNA or protein isoforms, although we did not detect any difference in mRNA levels for total p57 in the absence or presence of the paternal allele. The paternal allele thus also does not appear to affect the expression level of the maternal allele. It is also possible that transcripts other than p57 mRNA are produced from the paternal *p57* allele or from cis- or trans-regulatory elements within the paternal *p57* locus and contribute to the changes to the brain apparent in paternal p57 cKO mice.

Prenatal protein restriction and intrauterine growth retardation have been associated with various psychiatric conditions including schizophrenia, attention-deficit and hyperactivity disorder, autism spectrum disorder, and major affective depression^36^. Of interest in this regard, restriction of maternal dietary protein during pregnancy induces demethylation and aberrant induction of the *p57* locus in the prefrontal cortex and mesolimbic dopaminergic system of the resulting offspring^37^. Indeed, such protein restriction was found to result in permanent derepression of the imprinted paternal *p57* allele through a folate-dependent mechanism of DNA methylation loss^34^. However, whether or how such derepression of the paternal *p57* locus affects brain development remains to be clarified. Given that we found that the paternal *p57* allele plays an essential role in generation of appropriate numbers of NPCs and neurons, derepression of this allele induced by a low-protein diet may have an impact on NPC proliferation and neurogenesis during neocortical development. We also found that paternal p57 cKO resulted in a reduction in the staining intensity for tyrosine hydroxylase (TH) in the striatum (data not shown). Given that p57 has been implicated in the production of TH^+^ dopaminergic neurons^27^ and that twofold overexpression of p57 conferred by a transgene both increased TH expression in the brain and altered behaviors that are dependent on the mesolimbic dopaminergic system^38,39^, derepression of the paternal *p57* locus induced by a low-protein diet might thus lead to behavioral changes through modulation of neurogenesis in both the neocortex and mesolimbic dopaminergic system. The possible role of derepression of the paternal *p57* allele in the link between early-life adversity and aberrant brain development associated with psychiatric disorders thus warrants further investigation.

## Experimental Procedures

### Mice

C57BL6/J (BL6) mice were purchased from CLEA Japan. JF1/Ms (JF1) mice were provided by RIKEN BRC (RBRC00639) through the National BioResource Project of the MEXT/AMED, Japan, and purchased from National Institute of Genetics, Japan. Nestin-Cre mice and p57 floxed mice were kindly provided by Ryoichiro Kageyama and Keiichi Nakayama, respectively. To generate JF1 Nestin-Cre mice, Nestin-Cre mice were backcrossed to JF1 mice, and confirmed the SNPs at p57 genomic locus with sanger sequencing. All mice were maintained and studied according to protocols approved by the Animal Care and Use Committee of The University of Tokyo (approval numbers: P25-8, P25-27, PH27-3 and P30-4).

### FACS

Neocortex was prepared by manual dissection, digested enzymatically with a papain-based solution (Sumitomo Bakelite) and then resuspended in 0.3% BSA/PBS containing primary antibodies (PE-conjugated CD133 (mouse, 1:500, BioLegend, 141204) and APC-conjugated CD24 (mouse, 1:500, BioLegend, 101814)). Cells were then sorted with FACS Aria IIIu (BD). Debris and aggregated cells were gated out by forward and side scatter. Gating was done with isotype controls. NPCs and neurons were isolated as CD133^+^CD24^−^ fraction and CD133^−^ CD24^+^ fraction, respectively.

### RNA extraction and RT-qPCR

Allele specific qPCR was performed on cDNA prepared from sorted cells or neocortical tissues of F1 hybrid mice obtained by crossing BL6 and JF1. For other primer sets, qPCR was carried out with the use of cDNA prepared from neocortical tissues of BL6 mice. Total RNA was extracted using RNAiso Plus (Takara) following the instructions of the manufacturer. Reverse transcription (RT) was performed with at the maximum of 0.5 μg of total RNA and ReverTra Ace qPCR Master Mix with gDNA remover (TOYOBO). The obtained cDNA was subjected to real-time PCR analysis in a Roche LightCycler 480 II with THUNDERBIRD SYBR qPCR kit (TOYOBO). As for allele specific qPCR, plasmid DNA containing each SNPs was used as a standard and relative copy number was calculated according to the molecular weight. In other experiments, the amount of mRNA quantified was normalized relative to that of β-actin mRNA.

The used primers were as follows:

β-actin

Fw AATAGTCATTCCAAGTATCCATGAAA

Rv GCGACCATCCTCCTCTTAG

p21

Fw CACCTCTAAGGCCAGCTA

Rv AGCAATGTCAAGAGTCGG

p57

Common GGGCAGTACAGGAACCATTTC

BL6 TTAGCTTACAGTGTCCCGCA

JF1 TTAGCTTACAGTGTCCCGTA

Allele specific qPCR primers were designed as previously described^40^. BL6 /JF1 SNP sites (BL6 chr7: 143458462, NC_000073, GenBank) were identified using NIG Mouse Genome Database (http://molossinus.nig.ac.jp/msmdb/index.jsp). Total p57 mRNA was quantified with common and BL6 primers.

### Immunohistochemistry

For immunofluorescence staining, brains were postfixed with ice-cold 4% (wt/vol) paraformaldehyde (PFA) at 4 °C for 2 h, equilibrated with 30% (w/v) sucrose in PBS, and frozen in OCT (Tissue TEK). Coronal sections (15-16 μm thickness) were exposed to Tris-buffered saline (TBS) containing 0.1% Triton X-100 and 2% Donkey serum (blocking buffer) and incubated overnight at 4 °C with primary antibodies in blocking buffer and then for 1-2 h at room temperature with Alexa Fluor-conjugated secondary antibodies in blocking buffer. For staining with the antibody to Cux1, Ctip2, Pax6, Tbr2 and Ki67, we performed antigen retrieval by autoclave treatment of sections with target retrieval solutions (Dako) for 10min at 105°C. Fluorescence images were obtained with a laser confocal microscope (Leica TCS-SP5). Antibodies used for immunostaining included Cux1 (Rabbit, 1:200, Santa Cruz, sc-13024), Ctip2 (Rat, 1:1000, Abcam, ab18465), NeuN (Mouse, 1:200, Abcam, MAB377), S100β (Rabbit, 1:500, Abcam, AB52642) Pax6 (Rabbit, 1:1000, Millipore, AB2237), Tbr2 (Chicken, 1:1000, Millipore, AB15894), Ki67 (Rat, 1:500, Dako, M7249) cleaved Caspase3 (Rabbit, 1:1000, Cell Signaling, 9664). Alexa-Fluor-labeled secondary antibodies (1:1000) and Hoechst were obtained from Life Technologies.

### Statistical analysis

Data are presented as means + s.e.m. as indicated, and were analyzed by Student’s two-tailed paired *t* test and Student’s two-tailed unpaired *t* test, as indicated. A *P* value of <0.05 was considered statistically significant and the significance is marked by \**P*<0.05 and \*\**P*<0.01. The number of animals in each experiment is stated in the respective figure legends.

## Acknowledgments

We thank Toshihiko Shiroishi for providing us the genomic information of the JF1/Ms mouse strain. This study was supported by KAKENHI grants from the Ministry of Education, Culture, Sports, Science, and Technology of Japan and the Japan Society for the Promotion of Science (JP18K06477 for D.K.; JP16H06479, JP15H05773 for Y.G.) as well as by AMED-CREST of the Japan Agency for Medical Research and Development, by Uehara Foundation and by the International Research Center for Neurointelligence (WPI-IRCN) at The University of Tokyo Institutes for Advanced Study.

## Author contributions

Y.I. performed the experiments, analyzed the data, and wrote the manuscript. T.W. designed the allele specific qPCR of p57. H.M. conducted initial analysis of p57 paternal cKO mice. D.K., S.F. and Y.G. conceived of and coordinated the project as well as wrote the manuscript. All authors approved the final version of the manuscript.

---

**Figure S1**

**CNS-specific deletion of the p57 paternal allele reduced the thickness of upper layers**

**(a-b)** Cortical thickness of upper layer neurons (Cux1^+^) and deep layer neurons (Ctip2^+^) in the somatosensory area was quantified at P24 (**a**) and P0 (**b**)

Data are mean+s.e.m. Unpaired two-tailed Student’s t-test; n=3 mice/ group in (**a**), n=5 mice /group in (**b**); **\**P*** <0.05. n.s., not significant.

