## Supplementary figures and images for "Role of the imprinted allele of the p57^Kip2^ gene in mouse neocortical development"

### Supplementary Fig. 1

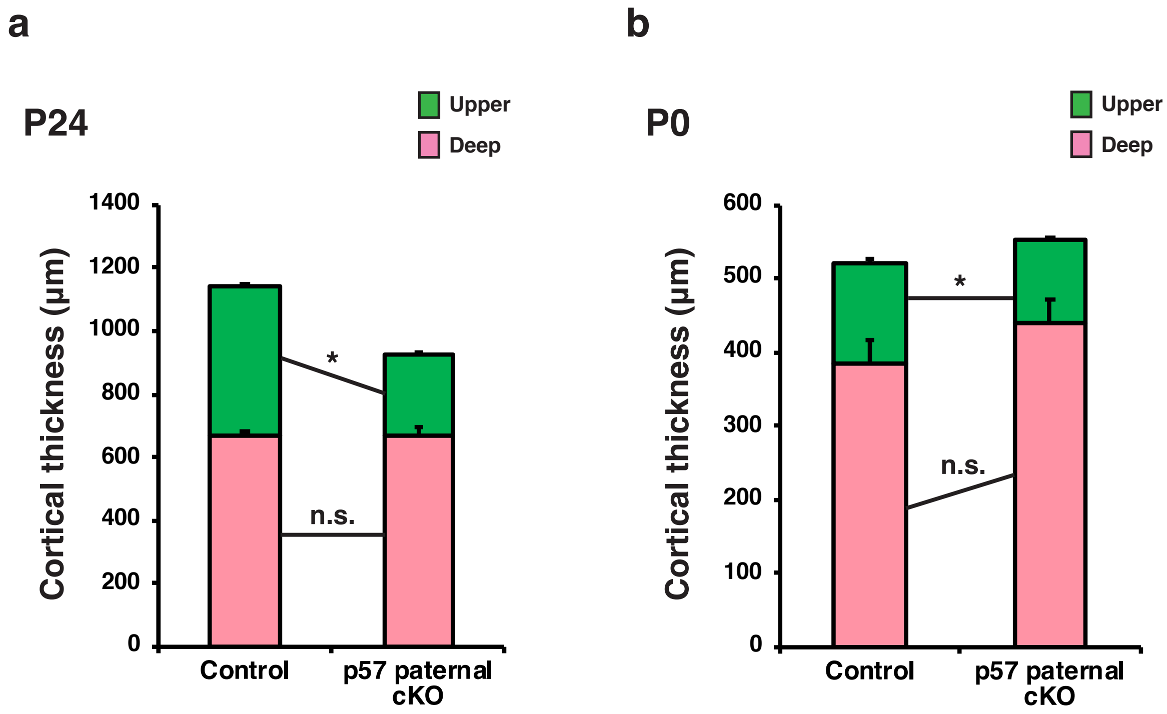
